# Applying environmental DNA metabarcoding to calculate an index of biotic integrity for freshwater fish

**DOI:** 10.1101/2025.05.26.656190

**Authors:** Amanda N. Curtis, Lynsey R. Harper, Mark A. Davis, Eric R. Larson

**Author notes:** Corresponding author: Eric R. Larson, Department of Natural Resources and Environmental Sciences, University of Illinois Urbana-Champaign, 1001 S. Goodwin Ave., Urbana, IL, United States of America.

## Abstract

Multi-metric indices like the Index of Biotic Integrity (IBI) are important biomonitoring tools for Clean Water Act compliance in the United States (US) but have also been implemented worldwide across many taxonomic groups. Environmental DNA (eDNA) metabarcoding could complement IBIs by increasing detection sensitivity for rare taxa while also reducing monitoring costs. To date, no studies have examined the efficacy of using eDNA metabarcoding to calculate IBIs in the US. Here, we used eDNA metabarcoding to calculate a fish-based IBI for streams and rivers of the Tennessee River Basin in northern Alabama, US. We collected water samples from 50 stream and river sites across a gradient of land use intensity, extracted eDNA from these samples, and sequenced the eDNA using vertebrate-specific primers. We compared our eDNA-IBI to a previous fish-IBI implemented by the state of Alabama using conventional sampling, as well as predicted biological condition of these streams from the US Environmental Protection Agency (EPA) based on a benthic macroinvertebrate multi-metric index (BMMI). We found a highly significant, positive relationship between the eDNA-IBI and fish-IBI for the same streams but a weaker, non-significant positive relationship between the eDNA-IBI and BMMI. Notably, the former was recovered despite eDNA-IBI sampling being conducted nine years after the conventional fish sampling at slightly different locations, the fish-IBI having used greater sampling effort throughout the year (including spring and autumn rather than only summer sampling), and a lack of reference DNA sequences that prevented eDNA detection for some species detected by conventional fish sampling. Accordingly, our study provides a baseline for how an eDNA-IBI may work relative to multi-metric indices calculated from conventional sampling, which can be improved through future directions identified and discussed in our paper.

## 1 INTRODUCTION

Freshwater organisms are disproportionally at risk of extinction compared to marine or terrestrial taxa (Ricciardi and Rasmussen, 1999; Burkhead, 2012). At least 39% of freshwater fish in North America are imperiled, a value that has increased since 1989, and freshwater fish extinction rates are estimated to be 877 times greater than background rates (Jelks et al., 2008; Burkhead, 2012). Human alterations to landscapes have resulted in freshwater biota declines due to changes in flow regimes, increased non-native species invasions, and runoff of pollutants, nutrients, and sediment (Dudgeon et al., 2006; Reid et al., 2019). Habitat degradation is the main cause of freshwater fish declines in North America, especially for species with narrow endemic ranges (Dudgeon et al., 2006; Jelks et al., 2008). In response to these threats to freshwater ecosystems, the United States (US) enacted federal legislation, most notably the Clean Water Act (CWA), to reduce discharge of pollutants and maintain fishable and swimmable waters. Initial enforcement of the CWA largely focused on chemical compliance and omitted the biological component (Karr et al., 1991). Because the CWA specifically includes biological integrity, biological communities also need to be monitored along with chemical water quality and physical habitat conditions (Karr, 1981; Karr et al., 2022).

In response to the need to measure biotic integrity, multi-metric indices such as the Index of Biotic Integrity (IBI) were developed (Yoder and Kulick, 2003; Kuehne et al., 2017). The first IBI globally was developed using Midwestern US fish communities and shown to be a useful measure of human influence on streams (Karr, 1981). Fish are good indicators because they are relatively easy to collect, identify, and vary by species in their tolerance to stressors like chemical pollution, sedimentation, and habitat loss or fragmentation (Fausch et al., 1984; Angermeier and Karr, 1986). The goal of an IBI is to rank habitats or ecosystems relative to minimally impacted reference conditions for that ecoregion by metrics including community composition (e.g., number of species, number of tolerant species) and individual organism health, providing a single measure of habitat or ecosystem quality across individual, population, community, and ecosystem levels (Karr, 1981; Karr, 1991). Since the initial fish-based IBIs for application to the CWA in the US, other biotic multi-metric indices have been developed globally and adapted for use with other taxa and ecosystems (e.g., Ruaro et al., 2020; Karr et al., 2022).

Although IBIs have been widely implemented, there are numerous limitations or criticisms in using these multi-metric indices to infer ecological conditions. For example, sampling methodology can affect species detections and resulting IBI scores (Angermeier and Karr, 1986), sampling can be costly and time-intensive (Evans et al., 2017), collection methods can harm organisms and cause mortality (Snyder, 2003), and this approach requires taxonomic expertise that is in decline (Kim and Byrne, 2006). However, the collection and identification of environmental DNA (eDNA) may improve or complement IBI methodologies (Baird and Hajibabaei, 2012; Keck et al., 2017; Pawlowski et al., 2021; Karr et al., 2022). eDNA is genetic material released by organisms into their environment, which can be extracted from environmental media (e.g., soil, water, or air) to detect species (Taberlet et al., 2012; Cristescu and Hebert, 2018). eDNA sampling can outperform conventional sampling in detecting rare or low abundance species and provide a more time- and cost-efficient alternative to conventional surveys (Smart et al., 2016; Fediajevaite et al., 2021, Sternhagen et al., 2024).

Advancements in eDNA analysis have progressed from detection of single target species to detection of whole communities through eDNA metabarcoding (e.g., Deiner et al., 2016; Pont et al., 2018). While omissions in reference DNA sequence libraries and primer biases can limit the performance of eDNA metabarcoding for some taxa (e.g., Wangensteen et al., 2018; Elbrecht et al., 2019), this approach has generally outperformed conventional sampling in detecting species across a broad suite of ecosystems (e.g., McElroy et al., 2020; Moss et al., 2022). Therefore, eDNA metabarcoding could complement or be incorporated into IBI biomonitoring (Pawlowski et al., 2018) for the following reasons: 1) increased detection sensitivity for rare, low abundance, or cryptic species (Fediajevaite et al., 2021); 2) increased species identification in regions with high diversity and/or with multiple morphologically similar species that would otherwise require taxonomic expertise (Becker et al., 2015; Fueyo et al., 2024); and 3) reduced overall cost and time of sampling allowing for an increased number of sampling sites (Evans et al., 2017). Further, eDNA metabarcoding has been successful in biomonitoring and calculating biotic indices for aquatic macroinvertebrates and diatoms (e.g., Pawlowski et al., 2018; Apothéloz-Perret-Gentil et al., 2021; Meyer et al., 2021; Suren et al., 2024), yet few studies to date have used eDNA metabarcoding to calculate a fish-based IBI (David et al., 2021; Yang et al., 2023) and to our knowledge none have been conducted in the US where fish IBIs are a major component of biological monitoring under laws like the CWA.

We calculated an eDNA-IBI for streams and rivers of the Tennessee River Basin in northern Alabama, US, based on the previous fish-based IBI of O’Neil and Shepard (2010). We compared our eDNA-IBI for a subset (58%) of our study sites to O’Neil and Shepard (2010)’s fish-IBI, who had sampled shared streams approximately a decade (nine years) before our study, as well as to model-predicted stream biological condition from the US Environmental Protection Agency (EPA) for our study sites (Fox et al., 2017; Hill et al., 2017). A successful eDNA-IBI would recover equivalent trends to O’Neil and Shepard (2010) or Hill et al. (2017), identifying poor biological condition (low IBI scores) or good biological condition (high IBI scores) at these same streams. We further investigated mechanisms behind eDNA-IBI performance relative to these other multi-metric indices by decomposing the eDNA-IBI into its component metrics for comparison to O’Neil and Shepard (2010), and exploring mechanisms behind non-detection of fish species anticipated to be present at our streams by eDNA like sequence coverage in the DNA reference library used. We conclude by highlighting potential advantages of using an eDNA-IBI, the pitfalls of such an approach, and how future studies may improve eDNA-IBIs.

## 2 METHODS

### 2.1 Study region

The Tennessee River Basin in northern Alabama, US (Figure 1), is renowned for its freshwater biodiversity in North America, including high endemism of crayfish, mussel, and fish species (Lydeard and Mayden, 1995). Over 170 fish species are believed to inhabit the river basin, many of which are rare, imperiled, and have limited distributions (Boschung Jr. and Mayden, 2004). This region has low coverage of protected areas despite its high aquatic diversity and threats from rapid human population growth (Elkins et al., 2019). A fish-based IBI specific to the Tennessee River Basin of Alabama was developed by O’Neil and Shepard (2010), which allowed us to adapt this IBI for use with eDNA metabarcoding data.

**FIGURE 1.**
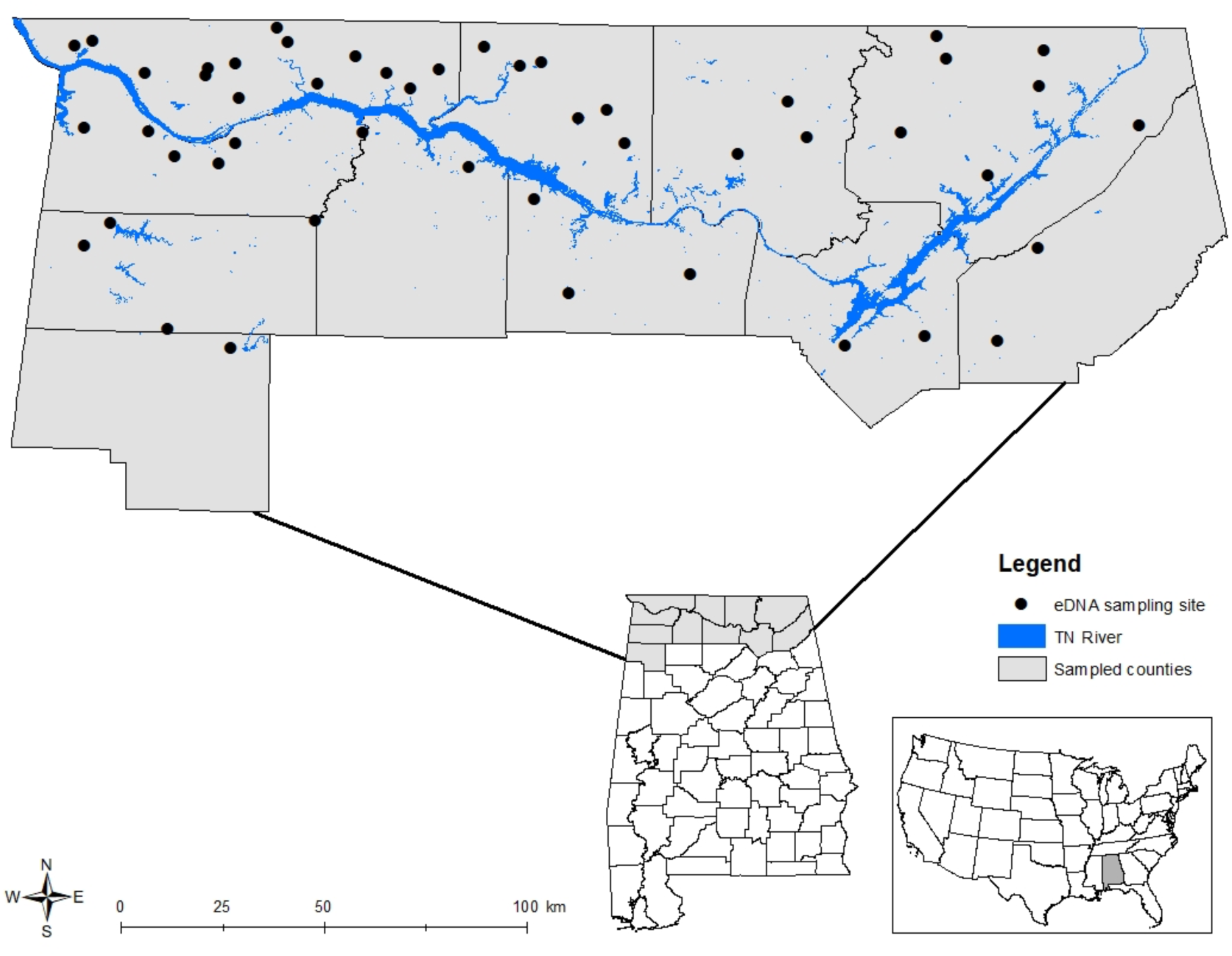
Stream sites (black dots) where water samples were collected for eDNA metabarcoding across the Tennessee River Basin (blue) in northern Alabama, US. Insets show the locations of sampled counties in northern Alabama and the location of Alabama in the US.

### 2.2 Environmental DNA sampling and extraction

From 25 July ‒ 11 August 2018, we sampled 50 stream and river sites across the Tennessee River Basin in Alabama (Figure 1). We selected sampling sites across a gradient of land use (rural, forested streams to highly urbanized streams) and stream size (4.08‒1122.5 km^2^ watershed area). We collected these water samples to monitor eastern hellbender salamanders (*Cryptobranchus alleganiensis alleganiensis*) with eDNA (Curtis, 2022) and repurposed the eDNA samples for this study. Repurposing of eDNA samples allowed us to examine the efficacy of an eDNA-IBI in a cost-effective manner with a limited budget and such repurposing has been successfully employed by others to ask multiple questions using archived samples (Dysthe et al., 2018; Franklin et al., 2019; Mori et al., 2023).

We collected four 250 mL surface water samples across the width of each stream site by wet-wading, with the exception of using a kayak at one deeper site (Elk River). We collected water upstream of the sampling individual, using the downstream flow of current to reduce risk of contamination, and took water samples only after suspended sediment from wading had settled. Polyethylene sample bottles (Thermo Scientific™ Nalgene™, Rochester, New York, US) had been previously washed in ∼50% v/v bleach solution (made from commercial Clorox containing ∼6% sodium hypochlorite), soaked in and triple rinsed with distilled water, dried, and bagged prior to use. At each site, we triple rinsed bottles with stream water prior to collecting eDNA samples. After collecting water samples, we sealed bottles in a bag and stored them on ice in a cooler until filtration at field housing. We used one field blank (250 mL) of distilled water per site to assess the potential for contamination in the field. Gloves were worn to collect all water samples and changed between sites. Prior to filtration of water samples, we decontaminated surfaces with bleach solution (∼50% v/v). All samples were filtered within 12 h of collection (Curtis et al., 2021a) onto cellulose nitrate filters (1.0 µm, 47 mm Whatman™, GE Healthcare, Chicago, Illinois, US). Sterile forceps were used to submerge filters into ∼1 mL of cetyltrimethylammonium bromide (CTAB), then filters were transported to the university laboratory, and frozen (−80°C) until DNA extraction. All water bottles, filter funnels, and forceps were washed in ∼50% v/v bleach solution, soaked in distilled water, and then triple rinsed with distilled water prior to use.

We extracted eDNA samples using a chloroform-isoamyl alcohol extraction procedure (Renshaw et al., 2015) and eluted to a final volume of 200 µL. We used one extraction blank consisting of extraction buffers and molecular grade water per every ∼12 samples to examine the potential for contamination in the lab. Samples were then analyzed for *C. a. alleganiensis* eDNA (Curtis, 2022) and frozen at −20°C until library preparation for metabarcoding. We cleaned work surfaces with a bleach solution (∼50% v/v), treated hoods with a UV cycle for 30 min, used a UV crosslinker on pipettes for at least 10 min, and frequently changed gloves to minimize contamination risk between eDNA samples during all laboratory procedures (extractions, PCR, pooling, etc.).

### 2.3 Environmental DNA metabarcoding

We tested existing vertebrate primer sets *in silico* (ecoPCR; Ficetola et al., 2010) against a custom database of DNA sequences for potential vertebrate species in northern Alabama collected from NCBI GenBank in March and April of 2020 (see Supplemental Information). We selected the 12S rRNA primers (12S-V5-F: 5′-ACTGGGATTAGATACCCC-3′ and 12S-V5-R: 5′-TAGAACAGGCTCCTCTAG-3′) developed by Riaz et al. (2011), as these primers amplified the greatest number of species *in silico*. We followed a similar two-step PCR library preparation procedure as Harper et al. (2019) and Di Muri et al. (2020). Each sub-library contained at least two PCR negative controls (molecular grade water) and two PCR positive controls of greater prairie chicken (*Tympanuchus cupido*), which is a bird native to the Midwestern US not found in Alabama (Ross et al., 2006). *In silico* testing confirmed that the 12S primers (Riaz et al., 2011) would amplify *T. cupido* DNA. We extracted DNA from *T. cupido* muscle tissue using a Qiagen DNeasy® Blood & Tissue Kit (Hilden, Germany) and then diluted tissue DNA to ∼0.05 ng/µL.

In the first PCR, we used 24 unique index combinations, including heterogeneity spacers, primers, and pre-adapters. We ran 25 µL PCR reactions with 12.5 µL of Q5® High-fidelity 2x master mix (New England BioLabs® Inc., MA, US), 0.5 µL bovine serum albumin (Thermo Fisher Scientific™, Waltham, MA, US), 1.5 µL of each primer (10 µM), 7 µL of molecular grade water, and 2 µL of eDNA. We used the following PCR conditions: initial denaturation at 98°C for 5 min followed by 35 cycles of 98°C for 10s, 58°C for 20s, 72°C for 30 s, and a final extension step at 72°C for 7 min. Reactions were prepared in strip tubes, with a drop of mineral oil added to each tube to prevent evaporation and minimize contamination risk (Di Muri et al., 2020). We prepared reactions and added eDNA in a dedicated clean room that was spatially separated from our high copy DNA laboratory and PCR machines. After eDNA was added and all tubes were sealed, we transported them to our main laboratory where we added our positive control. We performed PCR in triplicate with corresponding field and extraction blanks, in addition to PCR negative and positive controls. After PCR, we pooled sample/control replicates and visualized pooled PCR products on a 2% agarose gel. Sub-libraries were created by normalizing the PCR products based on the brightness of the PCR bands (5 µL for very bright, 10 µL for bright, 15 µL for faint bands, and 20 µL for very faint or no bands; Alberdi et al., 2018), 10 µL for each negative control (field, extraction, or PCR), and 1 µL for each positive control. To remove secondary bands, we used a double size selection magnetic bead (Mag-Bind® TotalPure NGS, Omega Bio-tek, Inc., Norcross, GA, US) clean-up procedure (Bronner et al., 2009) on each sub-library and confirmed absence of secondary bands by running sub-libraries on a 2% agarose gel.

In the second PCR, we attached the Illumina indexes to each sub-library in a 50 µL reaction comprised of: 25 µL Q5® High-fidelity 2x master mix (New England BioLabs® Inc., MA, US), 15 µL of molecular grade water, 3 µL of each primer (10 µm), and 4 µL of purified DNA. Reactions were prepared as described above, with sub-libraries run in duplicate. Sub-libraries were amplified using the following PCR conditions: initial denaturation of 95°C for 3 min, followed by 10 cycles of 98°C for 20 s and 72°C for 1 min, and a final extension at 72°C for 5 min. We pooled PCR replicates for each sub-library and visualized on a 2% agarose gel to confirm successful amplification. To remove secondary products, we again used a double size selection magnetic bead clean-up followed by confirmation of the absence of secondary bands on a 2% agarose gel. Sub-libraries were normalized based on number of samples they represented and estimated concentration using a Qubit™ dsDNA HS Assay Kit on a Qubit™ 3.0 fluorometer (Thermo Fisher Scientific™, Waltham, MA, US), then pooled for a final magnetic bead clean-up. Dilution and quantification of the final library concentration with real-time quantitative PCR using the KAPA library quantification kit (Roche, Wilmington, MA, US) on a CFX Connect Real-Time quantitative PCR system (Bio-Rad, Hercules, CA, US), library quality control using a Fragment Analyzer™ (Agilent Technologies, Santa Clara, CA, USA), and sequencing (library run at 10 pM with 10% PhiX Control v3 on an Illumina MiSeq using a MiSeq Reagent Kit v3 600-cycle) were performed by the Roy J. Carver Biotechnology Center, University of Illinois at Urbana-Champaign.

### 2.4 Bioinformatics analysis

We used a custom python script and the metaBEAT pipeline (v.0.97.11, https://github.com/HullUni-bioinformatics/metaBEAT) to conduct all bioinformatic processing (demultiplexing, quality trimming and merging, chimera removal, clustering, and taxonomic identification) following a similar methodology as Harper et al. (2019) and Moss et al. (2022), detailed in Supplemental Information. We used a minimum BLAST identity of 98% for taxonomic assignment of sequences relative to the custom vertebrate database. To account for potential false positive detections as a result of field or laboratory contamination, we applied sequence thresholds based on the maximum percentage of fish reads (0%) and non-fish reads (0.042%, white-footed mouse, *Peromyscus leucopus*) in any given PCR positive control (n = 32) to sequences for fish taxa and non-fish taxa in eDNA samples (see Supplemental Information for more detail). We also employed blank correction by removing the maximum number of reads for each taxon detected in negative process controls (n = 103) from reads for those taxa in eDNA samples. We then removed all singletons as potential sequencing artefacts, but retained all taxonomic assignments with ≥ 3 sequence reads to account for rare and potentially low-density taxa that may be underrepresented in sequencing. This refinement process accounts for potential false positive detections resulting from field or laboratory contamination as well as PCR or sequencing error whilst maximizing the number of fish species retained for eDNA-IBI calculation. Reads that could not be assigned to species level at 98% identity were removed from all subsequent analyses. Due to limited reference sequences and taxonomic debate around the genus *Campostoma* (Blum et al., 2008), matches to central stoneroller (*Campostoma anomalum*) and Mexican stoneroller (*Campostoma ornatum*) were manually reassigned to the largescale stoneroller (*Campostoma oligolepis*), as this is the only stoneroller species present in this region of Alabama. In rare instances where sequences were assigned to Japanese fluvial sculpin (*Cottus pollux*), we also manually reassigned to mottled sculpin (*Cottus bairdii*), as this was the only sculpin species detected in our samples and present in the Tennessee River basin of Alabama.

### 2.5 Index of Biotic Integrity calculation

We derived an eDNA-IBI from O’Neil and Shepard’s (2010) fish-IBI for the Tennessee River Basin of northern Alabama. From May ‒ October 2009, O’Neil and Shepard (2010) sampled 97 streams across a range of stream sizes and human stressor gradients, sampling each stream 32 times with a combination of backpack electroshocking and seining to target four different habitat types (i.e., 10 samples in riffles, 10 samples in runs, 10 samples in pools, and two samples at shorelines). O’Neil and Shepard (2010) initially considered 34 candidate metrics in developing their IBI. After examining relationships between these 34 metrics and habitat and human disturbance measures, 12 metrics were selected to calculate a combined IBI score (O’Neil and Shepard, 2010). These 12 metrics assessed species richness and composition, habitat disturbance, trophic structure, reproduction, and fish health to calculate an IBI score that O’Neil and Shepard (2010) considered successful in discriminating good from poor biological condition of streams and rivers in this region (Table 1).

**TABLE 1.**
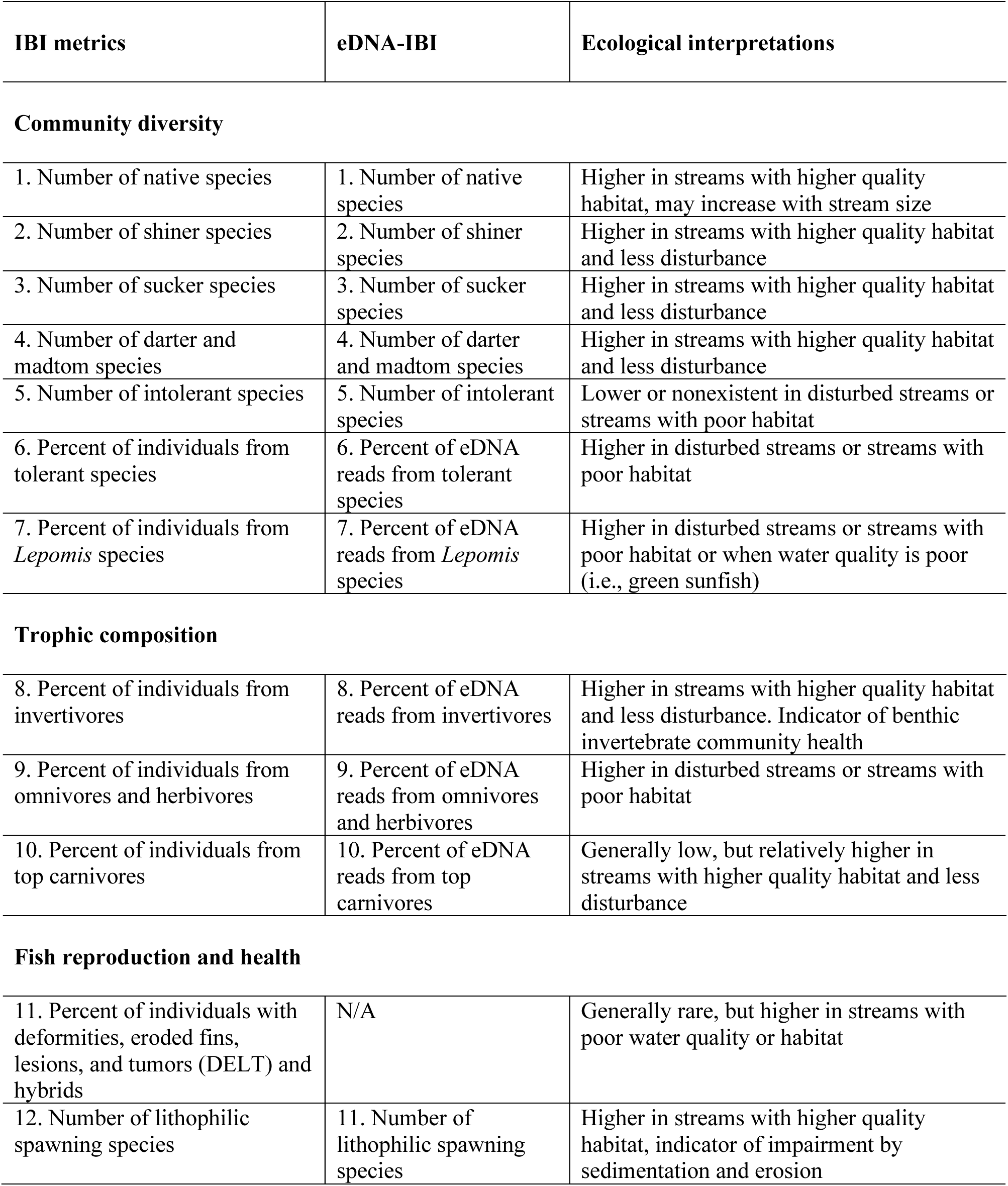
Index of biotic integrity (IBI) metrics from O’Neil and Shepard (2010) for freshwater fish of the Tennessee River Basin in northern Alabama with modifications for an environmental DNA IBI (eDNA-IBI) and ecological interpretations of IBI metrics.

| IBI metrics | eDNA-IBI | Ecological interpretations |
| --- | --- | --- |
| **Community diversity** |  |  |
| 1. Number of native species | 1. Number of native species | Higher in streams with higher quality habitat, may increase with stream size |
| 2. Number of shiner species | 2. Number of shiner species | Higher in streams with higher quality habitat and less disturbance |
| 3. Number of sucker species | 3. Number of sucker species | Higher in streams with higher quality habitat and less disturbance |
| 4. Number of darter and madtom species | 4. Number of darter and madtom species | Higher in streams with higher quality habitat and less disturbance |
| 5. Number of intolerant species | 5. Number of intolerant species | Lower or nonexistent in disturbed streams or streams with poor habitat |
| 6. Percent of individuals from tolerant species | 6. Percent of eDNA reads from tolerant species | Higher in disturbed streams or streams with poor habitat |
| 7. Percent of individuals from *Lepomis* species | 7. Percent of eDNA reads from *Lepomis* species | Higher in disturbed streams or streams with poor habitat or when water quality is poor (i.e., green sunfish) |
| **Trophic composition** |  |  |
| 8. Percent of individuals from invertivores | 8. Percent of eDNA reads from invertivores | Higher in streams with higher quality habitat and less disturbance. Indicator of benthic invertebrate community health |
| 9. Percent of individuals from omnivores and herbivores | 9. Percent of eDNA reads from omnivores and herbivores | Higher in disturbed streams or streams with poor habitat |
| 10. Percent of individuals from top carnivores | 10. Percent of eDNA reads from top carnivores | Generally low, but relatively higher in streams with higher quality habitat and less disturbance |
| **Fish reproduction and health** |  |  |
| 11. Percent of individuals with deformities, eroded fins, lesions, and tumors (DELT) and hybrids | N/A | Generally rare, but higher in streams with poor water quality or habitat |
| 12. Number of lithophilic spawning species | 11. Number of lithophilic spawning species | Higher in streams with higher quality habitat, indicator of impairment by sedimentation and erosion |

To calculate our eDNA-IBI, we first removed O’Neil and Shepard’s (2010) metric of fish health and hybridization (i.e., deformities, eroded fins, lesions, and tumors [DELT] or hybrids), as this information could not be obtained from our eDNA metabarcoding results (Table 1). Hybridization and DELT are rarely included in other IBIs (Karr et al., 1986; McCormick et al., 2001). Omission of this IBI metric results in the eDNA-IBI having a lower maximum value (55) than the fish-IBI (60) of O’Neil and Shepard (2010). Next, some IBI metrics of O’Neil and Shepard (2010) use species richness of native fishes or specific taxonomic groups, which is highly compatible with eDNA metabarcoding results, whereas other metrics use the percent of individuals collected at that site from pollution tolerant taxa (Table 2), including *Lepomis* sp. sunfishes, or particular trophic guilds (e.g., invertivore, omnivore, herbivore). In calculating the eDNA-IBI, we instead used relative read abundance from metabarcoding results to calculate percent-based metrics; for example, the percentage of reads from all tolerant species relative to the total number of reads at that study site. Contradictory results have been found on whether eDNA sequence reads can be used as a measure of fish abundance (Di Muri et al., 2020; Shelton et al., 2022), and we address in the discussion how this assumption may have influenced our results.

**TABLE 2.**
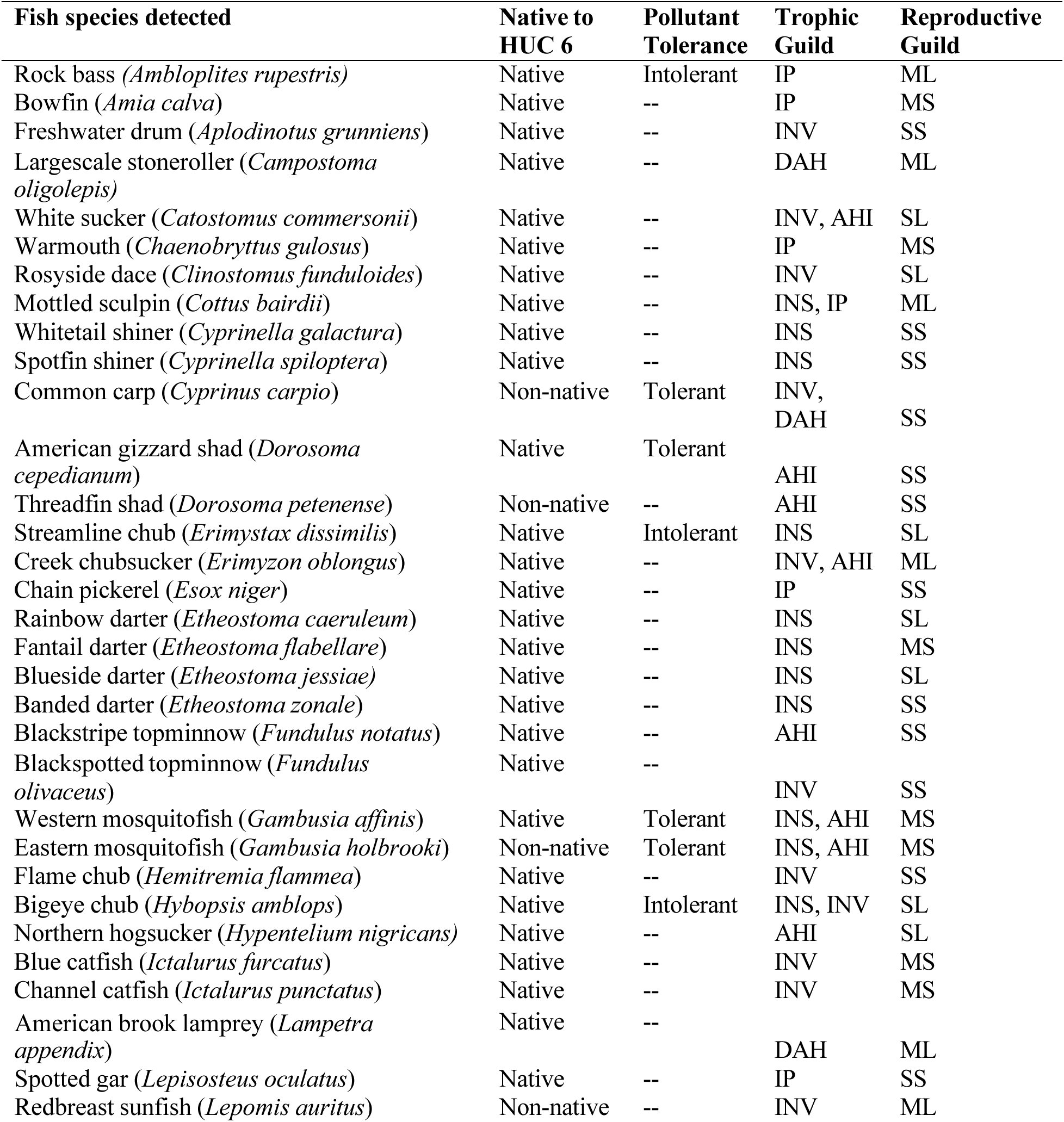
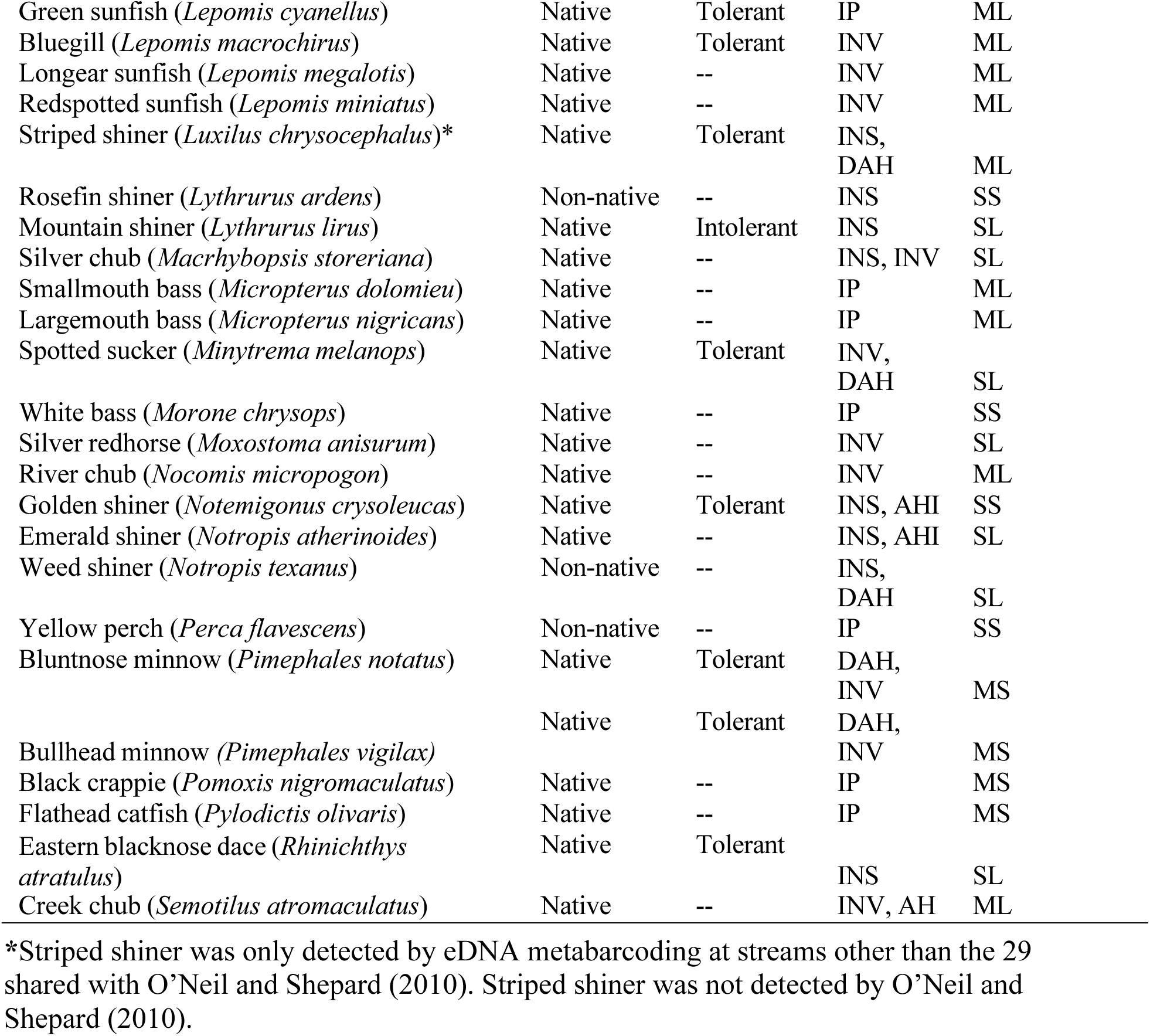
Fish species detected by environmental DNA metabarcoding in Tennessee River Basin streams of northern Alabama with associated information for calculation of index of biotic integrity metrics (Table 1). Native status from Boschung Jr. and Mayden (2004) and pollutant tolerance, trophic guild, and reproductive guild modified from O’Neil and Shepard (2010). Many species are intermediate between intolerant and tolerant to pollution (--). Trophic guilds include detritivore, algivore, herbivore (DAH); algivore, herbivore, invertivore (AHI); invertivore (INV); insectivore (INS); invertivore, piscivore (IP). Reproductive guilds include simple lithophils (SL); manipulative lithopils (ML); simple miscellaneous spawners (SS); manipulative miscellaneous spawners (MS).

We followed the methodology of O’Neil and Shepard (2010) to calculate numerical IBI scores. Watershed area or stream size can affect IBI scores (i.e., larger streams have higher species richness; Fausch et al., 1984), and accordingly we first examined relationships between each IBI metric and watershed area. For IBI metrics that showed evidence of a relationship with watershed area (number of darters + madtoms species, number of shiner species, number of native species), we used maximum species richness lines (includes 95% of the data) to trisect data into approximately equal divisions (Fausch et al., 1984) and assigned a 5-3-1 score, with a score of 5 as the best and 1 as the worst (O’Neil and Shepard, 2010). Most of our metrics (8 out of 11) showed no relationship with watershed area, so we followed O’Neil and Shepard (2010) in dividing data into approximate thirds scored as 5-3-1. As eDNA metabarcoding recovered different fish communities than O’Neil and Shepard’s (2010) conventional sampling (e.g., lower shiner species richness but higher species richness of intolerant taxa), different numerical thresholds are used to assign 5-3-1 scores for the same component metrics, emphasizing that these are similar but not identical multi-metric indices. Users of IBIs like O’Neil and Shepard (2010) often assign categorical integrity classes (e.g., very poor, poor, fair, good, excellent) to numerical IBI scores by percentiles (e.g., 10^th^, 25^th^, 50^th^, 75^th^, and 90^th^), but we have omitted reporting this step here to emphasize regression comparisons of numerical IBI scores between eDNA and conventional sampling (below).

### 2.6 Data analysis

We first compared the region-wide fish community (gamma diversity) recovered by eDNA metabarcoding to the community recovered by conventional sampling of O’Neil and Shepard (2010) for the 29 streams shared between studies. We next assessed site-level fish communities (alpha diversity) at these 29 streams by testing for differences in species richness using a paired Wilcoxon signed rank test with the *dplyr* package v1.1.0 (Wickham et al., 2023) in R v3.4.2 (R Core Team, 2022).

We regressed the fish-IBI scores of O’Neil and Shepard (2010) at the 29 shared streams against our eDNA-IBI scores. We sought to determine if the two multi-metric indices recovered statistically significant, shared relationships of biological condition at these streams, where lower values of the fish-IBI would correspond with lower values of the eDNA-IBI and higher values of the fish-IBI would correspond with higher values of the eDNA-IBI. We also regressed the quantitative values of the 11 shared metrics of both IBIs (Table 1), prior to scoring as 5-3-1, against each other to evaluate if there were metrics where eDNA metabarcoding performed well or poorly relative to conventional fish sampling.

Our comparison to O’Neil and Shepard (2010) was limited by the nine-year gap between our sampling events, although we anticipated little change in biological condition over our sampled gradient of anthropogenic stressors, from highly urban to remote, rural streams. Further, this comparison used 29 of our 50 sampled streams by eDNA, and we did not always sample the exact locations of O’Neil and Shepard (2010), differing by a mean distance of 3.04 km (<0.01 minimum to 16.00 km maximum distance; Table 3). We address implications of spatial and temporal mismatches between our eDNA metabarcoding and O’Neil and Shepard’s (2010) conventional fish sampling in the discussion. To use all our data with a better spatial match to stream biological condition, we regressed the benthic multi-metric index (BMMI) from US EPA (Fox et al. 2017; Hill et al., 2017) against our eDNA-IBI scores for our full dataset. The BMMI is a model-predicted probability (0 to 1) that a stream reach is in good biological condition based on the US EPA’s National Rivers and Streams Assessment for benthic macroinvertebrate communities (Hill et al., 2017). We downloaded BMMI data on 9 October 2020 from the US EPA StreamCat dataset (https://www.epa.gov/national-aquatic-resource-surveys/streamcat-dataset#access-streamcat-data; currently available at https://github.com/USEPA/StreamCatTools). All bivariate linear regression analyses (above) were conducted using the *lme4* package v1.1-32 (Bates et al., 2015) in R v4.1.0 (R Core Team, 2022).

**TABLE 3.** A summary of the 29 stream sites sampled by both O’Neil and Shepard (2010) and our study with distances (km) between sampling sites and fish species detected by either O’Neil and Shepard’s (2010) conventional sampling or environmental DNA (eDNA) metabarcoding (Figure 3).

| Stream | County | Distance between sites (km) | O’Neil and Shepard (2010) species | eDNA species |
| --- | --- | --- | --- | --- |
| Anderson Creek | Lauderdale | 5.08 | 11 | 5 |
| Bakers Creek | Morgan | 0.82 | 14 | 6 |
| Bear Creek | Marion | 0.05 | 9 | 8 |
| Big Coon Creek | Jackson | 0.08 | 24 | 10 |
| Bluewater Creek | Lauderdale | 5.26 | 26 | 13 |
| Bluff Creek | Lauderdale | 0.06 | 19 | 10 |
| Bumpass Creek | Lauderdale | 0.05 | 12 | 8 |
| Butler Creek | Lauderdale | 1.21 | 37 | 21 |
| Buzzard Roost Creek | Colbert | 8.84 | 18 | 5 |
| Cedar Creek | Franklin | 0.95 | 22 | 14 |
| Cox Creek | Anderson | 0.01 | 20 | 9 |
| First Creek | Lauderdale | 0.05 | 14 | 14 |
| Flat Rock Creek | Jackson | 0.01 | 8 | 6 |
| Flint River | Madison | 16.00 | 29 | 11 |
| Hurricane Creek | Jackson | 1.29 | 28 | 8 |
| Little Bear Creek | Franklin | 0.10 | 22 | 7 |
| Little Cypress Creek | Lauderdale | 7.09 | 26 | 21 |
| Muddy Fork | Lawrence | 3.89 | 16 | 9 |
| Paint Rock | Jackson | 3.03 | 32 | 11 |
| Pinhook Creek | Madison | 1.62 | 17 | 14 |
| Roseberry Creek | Jackson | 2.94 | 10 | 11 |
| Second Creek at Co. Rd 72 | Lauderdale | 0.04 | 16 | 11 |
| Second Creek at Ford’s Mill | Lauderdale | 0.16 | 27 | 10 |
| Short Creek | Marshall | 7.15 | 8 | 4 |
| South Sauty Creek | DeKalb | 0.08 | 11 | 6 |
| Sugar Creek | Limestone | 4.59 | 27 | 22 |
| Sulphur Creek | Limestone | 0.07 | 24 | 10 |
| Swan Creek | Limestone | 5.18 | 22 | 11 |
| Town Creek | DeKalb | 12.62 | 10 | 7 |

We did not recover some species detected by O’Neil and Shepard (2010) with eDNA metabarcoding and hypothesized that this might be due to lack of available 12S sequences on GenBank used to generate our custom vertebrate database, which can hinder taxonomic resolution to species-level (Jackman et al., 2021; Jerde et al., 2021). Alternatively, it is also possible that smaller-bodied or more benthic fish species might produce less eDNA or eDNA that is less detectable in the water column than larger-bodied or more pelagic fish species (Takeuchi et al., 2019; Andruszkiewicz Allan et al., 2021; Thalinger et al., 2021). Accordingly, we modeled detection of fish species by eDNA metabarcoding relative to O’Neil and Shepard (2010) for our shared 29 streams against the number of 12S sequences available for each species and their ecological traits. On 14 September 2022, we retrieved 12S data using the NCBI Web Search Tool (www.ncbi.nlm.nih.gov) and the NCBI Nucleotide Search incorporated into Geneious Prime v.2020.2.2 (https://www.geneious.com) to total the number of available 12S records for fish detected by either our eDNA metabarcoding or O’Neil and Shepard (2010), including complete mitogenomes. To match our species assignment in the bioinformatics pipeline, 12S records for *C. anomalum* and *C. ornatum* were used for northern Alabama’s *C. oligolepis.* For ecological traits (size and habitat), we accessed the FishTraits database (Frimpong and Angermeier, 2011) on 13 September 2022 to compile maximum total length (cm) for each fish species detected by either our eDNA metabarcoding or O’Neil and Shepard (2010). We ultimately omitted habitat traits, like benthic feeding, because most species in our dataset (85%) were benthic foragers per Frimpong and Angermeier (2011). We modeled eDNA detections against log transformed fish total length and log+1 transformed 12S sequences, due to zero values, with logistic regression in base R v3.4.2 (R Core Team, 2022).

## 3 RESULTS

The MiSeq run produced 30,014,808 reads. After bioinformatic filtering, 11,372,686 reads were retained, of which 8,991,559 reads (79.1%) were assigned to species-level and 4,894,629 to fish species (43.0%). The number of fish reads per stream ranged from 1,764 (Crow Creek) to 348,222 (Butler Creek). Overall, evidence of contamination or tag-jumping of fish species was generally low, but one extraction blank (Sep06.EB.03) contained sequences of largescale stoneroller (*Campostoma anomalum*, corrected to *Campostoma oligolepis*; 32 reads), rosyside dace (*Clinostomus funduloides*; 14 reads), bluegill (*Lepomis macrochirus*; 16 reads), striped shiner (*Luxilus chrysocephalus*; 14 reads), *Etheostoma* (28 reads), *Ictalurus* (15 reads), *Percina* (3 reads), and Percidae (21 reads). This contamination was accounted for by blank correction (see Methods and Supplemental Information). No fish sequences were found in any other negative process controls (see Supplemental Information).

Across the 50 streams sampled, we detected 56 fish species by eDNA metabarcoding. The most commonly detected species were largescale stoneroller (*C. oligolepis*), northern hogsucker (*Hypentelium nigricans*), striped shiner (*L. chrysocephalus*), largemouth bass (*Micropterus nigricans*), bluegill (*L. macrochirus*), and green sunfish (*Lepomis cyanellus*). We detected 4‒23 fish species per stream, with Short Creek, Elk River, and Town Creek having the fewest species detected and Cypress Creek having the most (Table 3). We detected 36 fish species in common with O’Neil and Shepard (2010) at the 29 streams sampled by both studies (Figure 2). O’Neil and Shepard (2010) detected 33 fish species that we did not detect, but we detected 19 fish species that O’Neil and Shepard (2010) did not detect. We detected only one species with eDNA metabarcoding, striped shiner (*L. chrysocephalus*), that was not recorded in Alabama by O’Neil and Shepard (2010); this was identified among samples from the 21 streams that only we sampled (Table 2). eDNA metabarcoding tended to underrepresent some small benthic species like darters and madtoms, but recovered two rare and narrowly distributed species missed by O’Neil and Shepard (2010): American brook lamprey (*Lampetra appendix*) in Bumpass Creek and flame chub (*Hemitremia flammea*) in Brier Fork of the Flint River, Cypress Creek, Piney Creek, and Swan Creek. However, at individual streams, fish species richness was significantly higher in O’Neil and Shepard (2010) than recovered by eDNA metabarcoding (V = 8.50, P < 0.001; Table 3; Figure 3).

**FIGURE 2.**
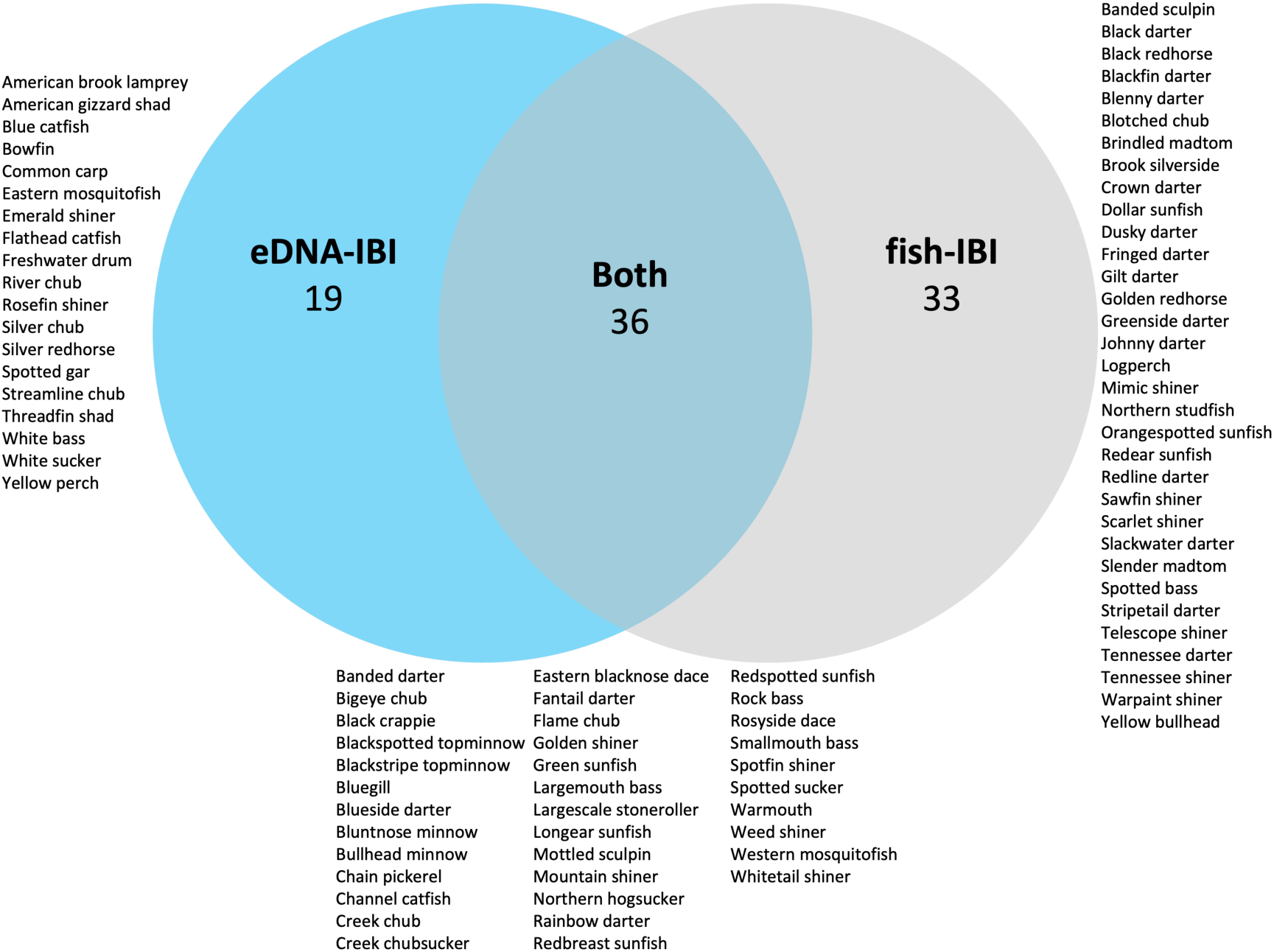
Fish species detected at the 29 stream sites sampled by both our study and O’Neil and Shepard (2010; Table 3) as species detected by both studies (intersection), O’Neil and Shepard (2010) only (fish-IBI: gray), or eDNA only (eDNA-IBI; blue).

**Figure 3.**
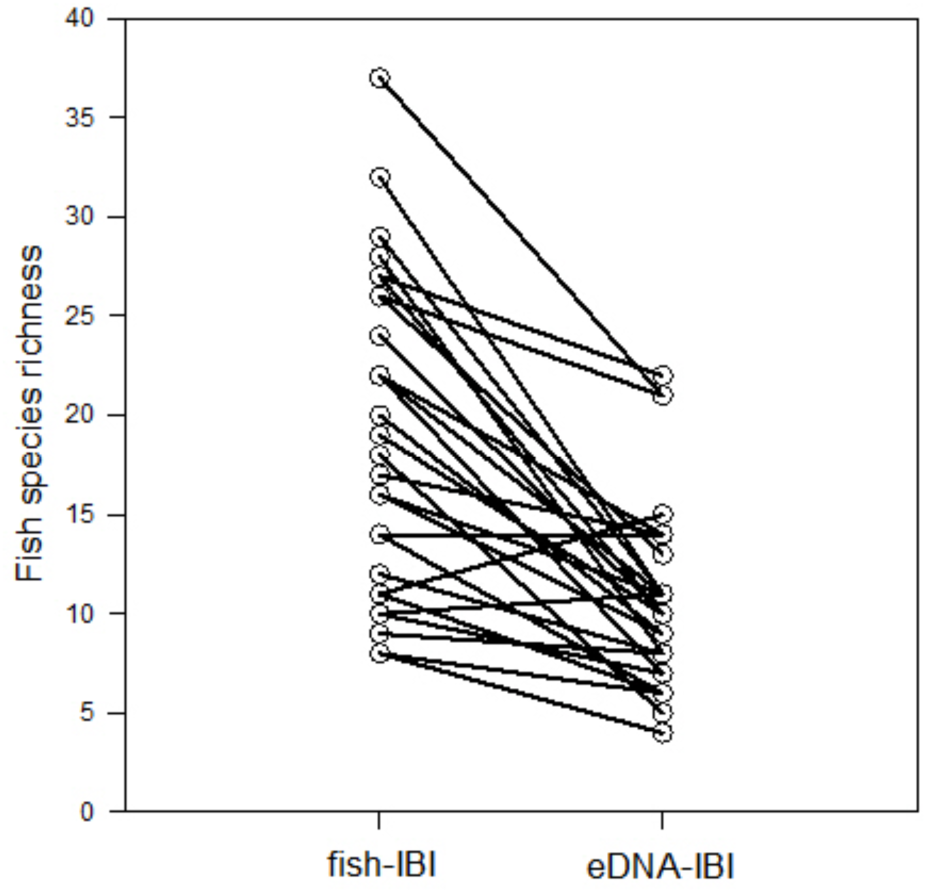
Fish species richness detected by O’Neil and Shepard’s (2010) fish-IBI relative to our eDNA-IBI at 29 streams sampled by both studies (Table 3).

Our eDNA-IBI scores across all 50 streams ranged between 19 and 43 with a mean score of 30.44. We found a positive, highly significant relationship between the fish-IBI and eDNA-IBI (R^2^ = 0.312, P = 0.002, Figure 4A), and a positive but weak and non-significant relationship between BMMI and eDNA-IBI (R^2^ = 0.064, P = 0.079, Figure 4B). We found six of 11 IBI metrics had significant relationships between the fish-IBI and eDNA-IBI, including the total number of species, number of darters+ madtom species, number of intolerant species, percent individuals as tolerant species, percent from *Lepomis* species, and number of lithophilic spawner species (Figure 5). The remaining five IBI metrics had poor relationships between the fish-IBI and eDNA-IBI, including number of sucker species, number of shiner species, percent from invertivore species, percent from omnivores+herbivores, and percent as top carnivores (Figure 5).

**FIGURE 4.**
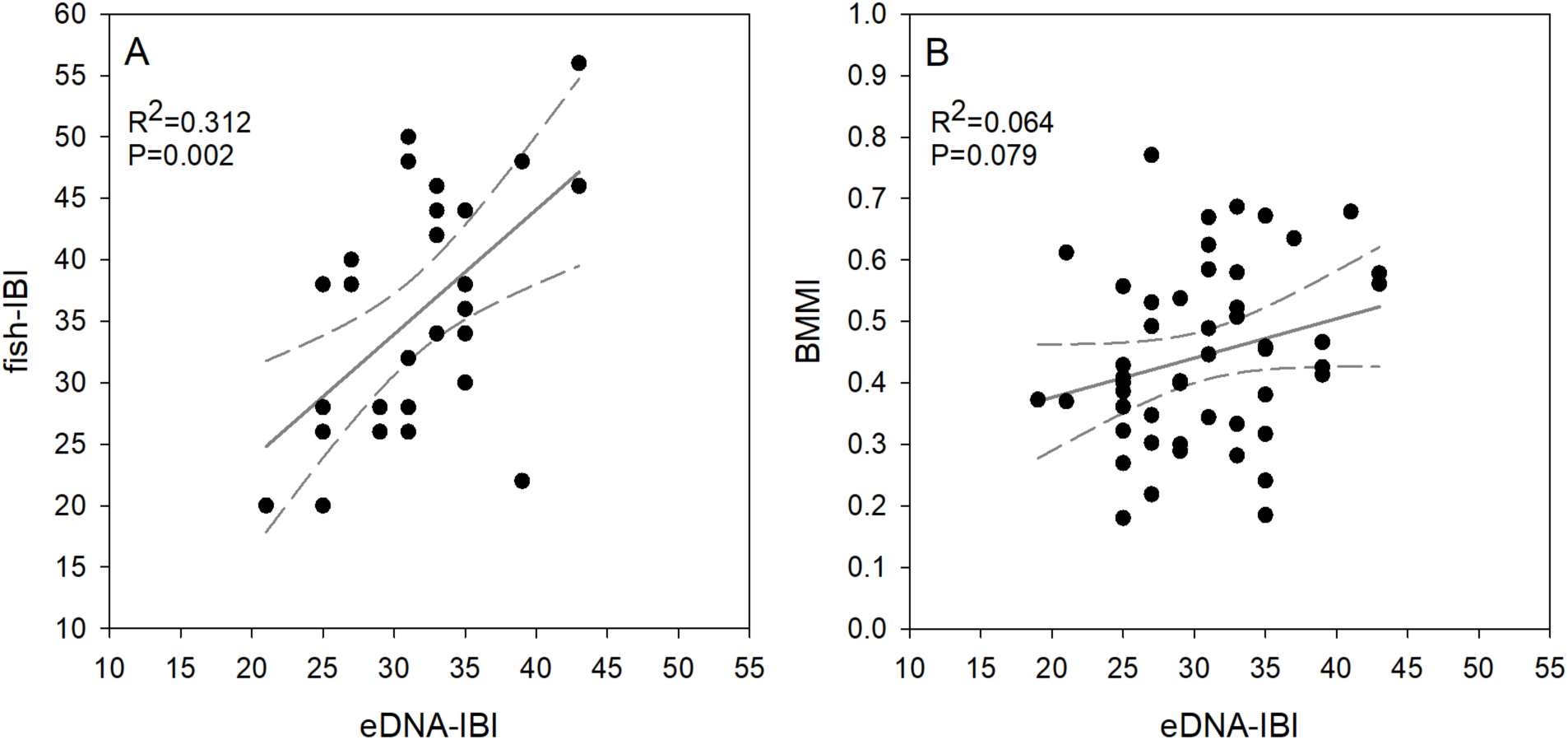
Bivariate regressions of the fish-IBI (A) of O’Neil and Shepard (2010) against our eDNA-IBI for 29 shared stream sites (Table 3) and the BMMI (B) of Hill et al. (2017) against our eDNA-IBI for the broader 50 stream sites we sampled (Figure 1). Axes are scaled to maximum and minimum possible values of the fish-IBI, eDNA-IBI, and BMMI, where the fish-IBI has a higher maximum value than the eDNA-IBI because it includes an additional metric that could not be calculated from eDNA data (Table 1). Linear regression lines (gray) are shown with 95% confidence intervals (gray dash).

**FIGURE 5.**
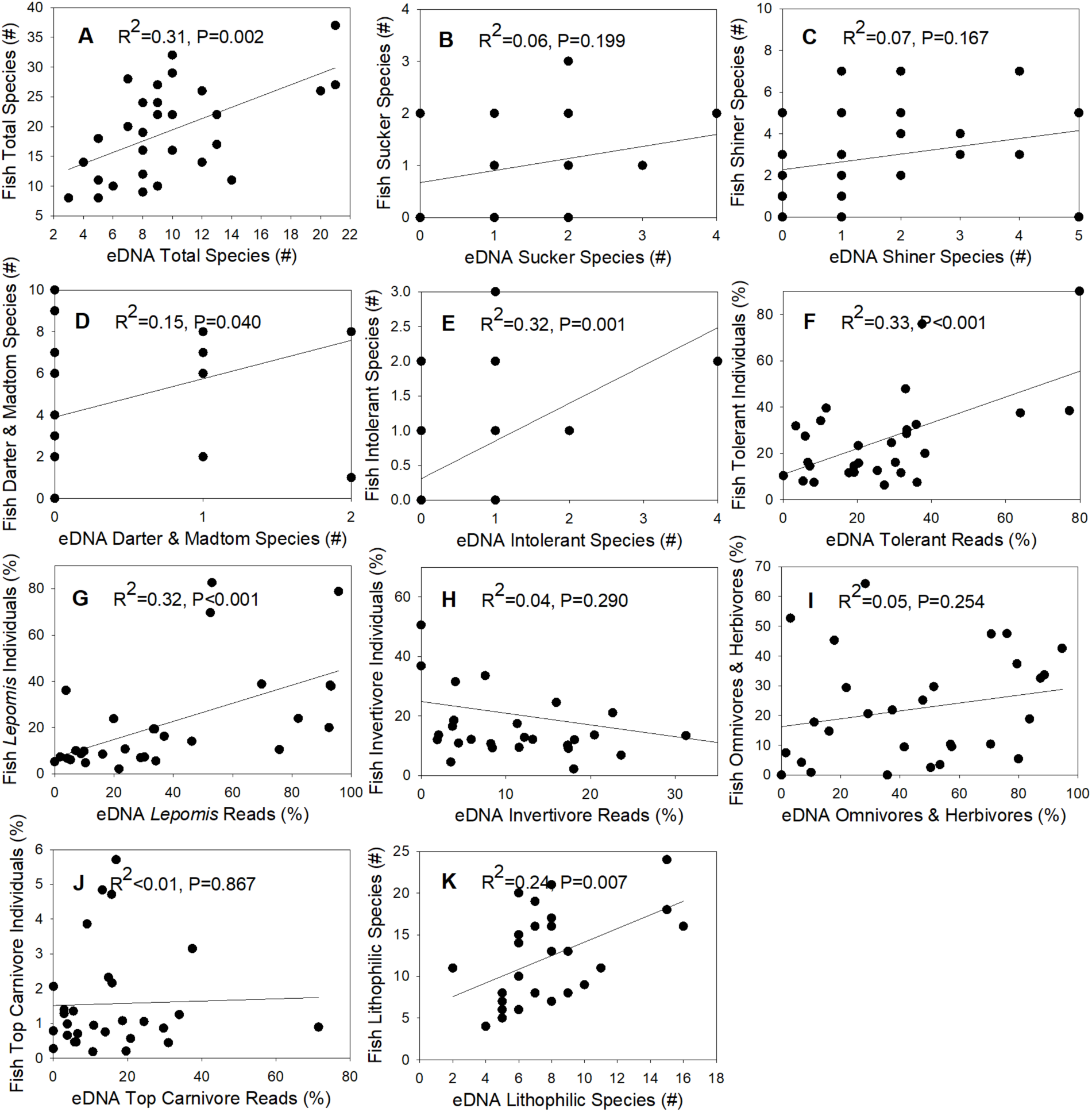
Relationships between 11 shared metrics of the fish-IBI from O’Neil and Sheperd (2010) and our eDNA-IBI from the 29 streams sites sampled in both studies (Table 3). (A) total number of fish species; (B) number of sucker species; (C) number of shiner species; (D) number of darters + madtom species; (E) number of intolerant species; (F) percent of individuals or reads from tolerant species; (G) percent of individuals or reads from *Lepomis* species; (H) percent of individuals or reads from invertivore species; (I) percent of individuals or reads from omnivore + herbivore species; (J) percent of individuals or reads from top carnivore species; (K) number of lithophilic spawning species. Linear regression relationships (gray) are shown without 95% confidence intervals for simplicity on this complex figure.

The number of 12S sequences available from GenBank (Coefficient ± SE = 4.679 ± 1.060, P < 0.001; Figure 6A) and maximum total length of fish (Coefficient ± SE = 1.249 ± 0.865, P = 0.149; Figure 6B) were both positively associated with detections in our eDNA metabarcoding data set, but this effect was only significant for sequence availability. Of 33 fish species detected by O’Neil and Shepard (2010) but not detected by eDNA metabarcoding, 14 (42.4%) lacked any 12S sequence data. Sequence availability was positively correlated with fish body size (Pearson R = 0.397, P < 0.001), but with low collinearity in our combined regression model (Variance Inflation Factor = 1.187).

**FIGURE 6.**
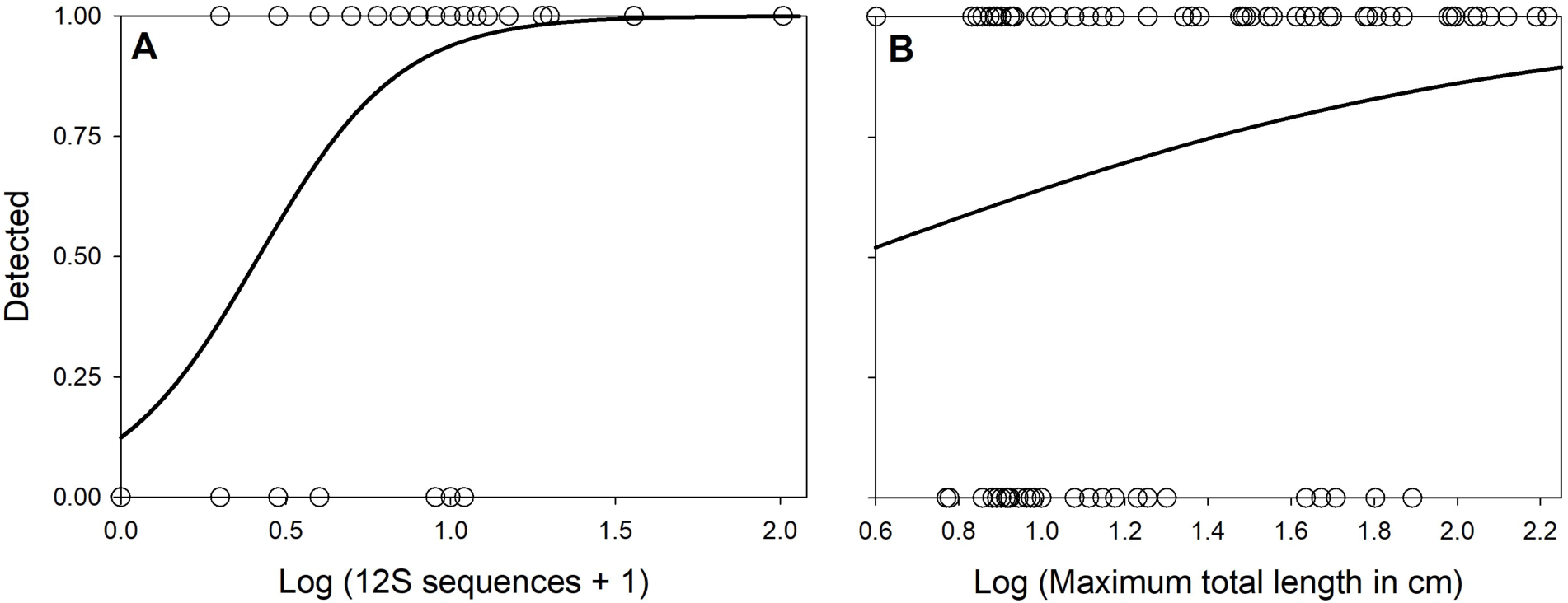
Relationships between the number of available 12S sequences (A) and maximum total length (B) of fish species and whether they were detected by eDNA metabarcoding at the 29 streams sampled by both our study and O’Neil and Shepard (2010; Figure 2).

## 4 DISCUSSION

We found that an eDNA-IBI and fish-IBI from conventional sampling recovered consistent, significant relationships of biological condition across 29 streams of northern Alabama, and this relationship was stronger than a comparison between the eDNA-IBI and model-predicted biological condition from benthic macroinvertebrate communities for a larger, 50 stream dataset. Our eDNA-IBI performed similarly to the fish-IBI even though eDNA metabarcoding detected fewer fish species overall, likely due to DNA reference library omissions and lower eDNA sampling effort (i.e., no spring or autumn sampling), and our sampling efforts were not perfectly aligned in space or time for these streams. This study establishes baseline performance for an eDNA-IBI in the US that can only improve in the future as DNA sequence libraries improve and through better alignment of eDNA and conventional sampling in space and time.

Environmental DNA metabarcoding failed to detect 33 species found by O’Neil and Shepard (2010) across our 29 shared streams, and detected significantly fewer fish species per stream. Notably, almost half of the species that were undetected by eDNA metabarcoding relative to the conventional fish sampling of O’Neil and Shepard (2010) lacked any 12S sequence data at GenBank, from which we derived our Alabama-specific reference database, and accordingly could not be detected by eDNA metabarcoding. Although total length was a weak predictor of eDNA detections relative to O’Neil Shepard (2010) when compared against 12S sequence coverage, total length was also significantly, positively correlated with availability of 12S sequences for these fish. Many small or benthic fish like the blackfin darter (*Ethoestoma nigripinne*), fringed darter (*Etheostoma crossopterum*), and scarlet shiner (*Lythrurus fasciolaris*) lacked 12S sequence data at the time of our analysis and were, as such, undetected by eDNA metabarcoding. Resolving deficiencies in DNA sequence libraries for many taxa remains a priority and would benefit the application of eDNA metabarcoding to freshwater bioassessment (Jerde et al., 2021; Marques et al., 2021).

Non-detections of fish species by eDNA metabarcoding with available 12S sequence data relative to O’Neil and Shepard (2010) could be due to several factors, including primer bias and competition for amplification with more common species (Nichols et al., 2018; Thalinger et al., 2021) as well as spatial and temporal mismatches between our two studies. For example, eDNA metabarcoding samples were collected in 2018 whereas conventional sampling by O’Neil and Shepard (2010) occurred in 2009. Accordingly, some fish species detected by O’Neil and Shepard (2010) may not have been present in streams when we sampled or may not have occurred at our slightly different sampling locations. We attempted to find and use fish-IBI data collected more recently relative to our eDNA sampling, but these data were too sparse for inclusion. Further, O’Neil and Shepard (2010) had higher sampling effort, through more of the year (May to October), than our study. We collected only four water samples (1 L of water total) per site, exclusively in late summer. This level of sampling effort is low relative to some other eDNA metabarcoding studies (e.g., Cantera et al., 2019; Di Muri et al., 2020). Seasonality of reproduction or migration timing can have a strong influence on the amount of eDNA present and detectability (Uchii et al., 2017; Curtis et al. 2021b). Our eDNA samples were initially collected to target late summer reproduction of a salamander species (Curtis, 2022), and may have failed to detect fish like the blenny darter (*Etheostoma blennius*) or brindled madtom (*Noturus miurus*) that spawn in the spring or early summer (Burr, 1979; Burr and Mayden, 1982). Future attempts to calculate an eDNA-IBI may benefit from collecting more field replicates or filtering larger volumes of water, sampling through more of the year, and sampling concurrently with conventional fish surveys to better facilitate comparisons between these methods (Curtis et al., 2021b; Macher et al., 2021).

Alternatively, eDNA metabarcoding detected 19 species that were not recorded by O’Neil and Shepard (2010) at the same streams, including rare species with limited distributions in northern Alabama as well as non-native species. For example, *H. flammea* is a Tennessee River Basin endemic that has been experiencing population declines and is considered vulnerable by the state of Alabama (Stallsmith, 2010). Stallsmith (2010) conducted a survey for *H. flammea* in known historical streams and found them abundant only in Cypress Creek and the Flint River. Our results support the presence of *H. flammea* in the Brier Fork of the Flint River, Cypress Creek, Piney Creek, and Swan Creek. Additionally, eDNA metabarcoding recovered some non-native fish species not detected by O’Neil and Shepard (2010), including threadfin shad (*Dorosoma petenense*), common carp (*Cyprinus carpio*), eastern mosquitofish (*Gambusia holbrooki*), and yellow perch (*Perca flavescens*). Nearly a decade separated our sampling from O’Neil and Shepard (2010), and these non-native species may have spread and/or become more common in northern Alabama since 2008. Alternatively, some of these non-native fish species may have been less detectable by O’Neil and Shepard’s (2010) conventional sampling methods due to low abundance (Jerde et al., 2011). A benefit of eDNA metabarcoding is that it can be repurposed for multiple, simultaneous environmental monitoring needs like invasive species surveillance (Dysthe et al., 2018; but see Darling et al., 2020).

Despite recovering slightly different fish communities by eDNA metabarcoding relative to conventional sampling, we found a positive, highly significant relationship between the eDNA-IBI and the fish-IBI at the same streams. This positive, highly significant relationship may not be surprising, as multi-metric indices like IBIs use functional traits, trophic guilds, and relative community composition to calibrate and assign biological condition to habitats and sites (Karr, 1981; Karr, 1991). Different field sampling methods do not have to recover identical communities to arrive at similar assignments of biological condition by multi-metric indices like IBI, especially over large gradients of anthropogenic stressors (Detenbeck and Cincotta, 2008; Vondracek et al., 2014). Alternatively, we recovered a positive but non-significant relationship between the eDNA-IBI and the US EPA’s BMMI at our full 50 stream dataset (Hill et al., 2017). The weaker relationship between the eDNA-IBI and BMMI may be surprising, as the BMMI values were from the same stream reaches as eDNA metabarcoding whereas the fish-IBI was occasionally mismatched in space. However, the eDNA-IBI may be better calibrated relative to the fish-IBI that it was closely derived from, or multi-metric indices based on different taxonomic groups may document different information about the condition of streams and rivers. For example, fish may be more responsive to physical habitat degradation and invertebrates more responsive to water quality (e.g., Pilière et al., 2014).

Our eDNA-IBI was closely derived from O’Neil and Shepard’s (2010) fish-IBI, but future development of eDNA-IBIs does not have to follow the same metrics that work best from conventional sampling. For example, we found significant, positive relationships between the eDNA-IBI and fish-IBI for total number of native species, number of darters + madtom species, number of intolerant species, number of lithophilic spawner species, percent as tolerant species, and percent as *Lepomis* species, but non-significant relationships for the number of sucker species, number of shiner species, percent from invertivore species, percent from omnivores+herbivores, and percent as top carnivores. As eDNA will work differently from conventional sampling for many taxonomic groups, and recover slightly different communities with different spatial or temporal contexts, multi-metric indices using eDNA should likely be calibrated anew following processes like O’Neil and Shepard (2010) and originating work like Karr (1981). Here, however, we establish that applying a conventional IBI to eDNA data with minimal modification recovers consistent relationships of stream biological condition relative to the originating IBI.

Environmental DNA metabarcoding has several strengths for use in freshwater bioassessment and biomonitoring, as well as weaknesses that may be improved by future methodological advances. First, several metrics of the IBI we used are calculated from percentages of fish relative abundance at their sampling site derived from conventional sampling. For consistency, we similarly used relative read abundance for those metrics, despite studies that question the use of reads as proxies for abundance (Elbrecht and Leese, 2015; Piñol et al., 2015). Conversely, other studies have found positive relationships between eDNA metabarcoding reads and fish relative abundance or biomass (Evans et al., 2016; Pont et al., 2018; Di Muri et al., 2020; Brantschen and Altermatt, 2024), and ongoing method development may strengthen links between metabarcoding reads and relative abundance for sampled communities (Ushio et al., 2018; Yates et al., 2021a). For example, estimates of relative abundance from eDNA metabarcoding reads may benefit from corrections for dilution by streamflow in lotic ecosystems (e.g., Curtis et al., 2021b) as well as differences in shedding rates between species based on laboratory studies (Andruszkiewicz Allan et al., 2021) or allometric scaling (Yates et al., 2022). The need for relative abundance from eDNA metabarcoding data could also be avoided by using observed-to-expected (O/E) predictive indices from presence/absence data rather than IBI (Hawkins et al., 2000), which is common in many biomonitoring applications for freshwater ecosystems (Kuehne et al., 2017; Comte et al., 2022). Alternatively, use of a taxon-independent community index, which does not require reads be assigned to species level but instead uses amplicon sequence variants (ASVs), could be explored as a way to estimate biological communities and infer ecological health (Wilkinson et al., 2024).

Second, we could not calculate an IBI metric from O’Neil and Shepard (2010) related to fish health and hybridization because this information was not available from our eDNA metabarcoding methodology. However, future use of CRISPR-Cas or nuclear DNA markers may allow eDNA-based estimates of hybridization (Williams et al., 2019; Stewart and Taylor, 2020; but see Couton et al., 2023), whereas environmental RNA (eRNA) may allow quantification of fish health or gene expression related to stress (Cristescu, 2019; Veilleux et al., 2021; Yates et al., 2021b). Lastly, a multi-locus approach with additional primers could be used to amplify known fish pathogens and/or parasites to provide measures of fish health (e.g., Peters et al., 2018; Katz et al., 2023), whereas human or livestock eDNA recovered from eDNA metabarcoding could be incorporated into freshwater bioassessment as possible indicators of agricultural impacts or wastewater discharge from point sources. Environmental DNA analysis, and related approaches like eRNA analysis, offer promising pathways to modify conventional bioassessment tools like IBI to simultaneously measure both stressors and population or community responses from the same environmental samples (Karr et al., 2022).

## 5 CONCLUSIONS

We found that eDNA metabarcoding recovered similar assessments of stream biological condition relative to conventional sampling of freshwater fish when used in the calculation of a multi-metric index, the IBI. Positive, highly significant relationships between the eDNA-IBI and a fish-IBI were recovered even though eDNA metabarcoding detected fewer fish species at both the regional and stream scale relative to conventional sampling. Improving molecular sequence coverage of freshwater fish from species-rich regions like the southeastern US should be a priority for improving integration of eDNA metabarcoding into bioassessment approaches like IBI. Further, some metrics of the IBI were significantly and positively associated between eDNA and conventional fish sampling data while others were not, and this information might be used to define and calibrate new, eDNA-specific multi-metric indices like the IBI. We are optimistic that ongoing eDNA research and development will likely improve estimates of organism abundance within communities (Yates et al., 2021a), characterize organism health or hybridization (Cristescu, 2019; Veilleux et al., 2021), and potentially identify the simultaneous presence of stressors like disease (Katz et al., 2023), enabling novel applications of eDNA to biological monitoring. However, our study establishes a baseline expectation that an IBI derived from eDNA survey should be highly similar to one derived from conventional sampling for freshwater fish.

## AUTHOR CONTRIBUTIONS

**ANC:** Conceptualization, Methodology, Analysis, Writing; **LRH:** Methodology, Analysis, Writing; **MAD:** Methodology, Analysis, Writing; **ERL:** Conceptualization, Methodology, Analysis, Writing

## ACKNOWLEDGEMENTS

We thank The Nature Conservancy in Alabama and Steve Northcutt for providing field housing. We are grateful to George Balto for fieldwork assistance and Cal Johnson, Rebecca Bearden, and Nathan Shrum for sharing fish data and conversations about Alabama fish communities. This manuscript was improved by comments from Robert L. Schooley, Michael P. Ward, and anonymous reviewers.

## DATA ACCESSIBILITY

Data associated with this study have been uploaded to the Illinois Data Bank, accessible at https://doi.org/10.13012/B2IDB-1381857_V1. Raw sequences from this study will also be uploaded to the NCBI Sequence Read Archive upon manuscript acceptance.

