## Supplemental Information for "Applying environmental DNA metabarcoding to calculate an index of biotic integrity for freshwater fish"

### 1 CUSTOM REFERENCE DATABASE

Following similar methodology as Harper et al. (2019) and Moss et al. (2022), custom reference databases were created for the 12S and 16S ribosomal RNA (rRNA) genes across all vertebrate species (including domestic animals) that could potentially occur within Alabama. Species checklists, information from the state of Alabama (www.outdooralabama.com), and Boschung Jr. and Mayden (2004) for fish species were used to compile species lists to query and download all available 12S and 16S rRNA sequences from the NCBI nucleotide database (https://www.ncbi.nlm.nih.gov/nucleotide). Our species list included 628 species composed of 84 herptile species, 163 fish species, 307 bird species, and 74 mammal species. Of these, 12S records were available for 410 species and reference sequences were missing for 218 species. Our 16S reference database included records for 367 species and reference sequences were missing for 261 species. Existing 12S (Cooper, 1994; Kitano et al., 2007; Riaz et al., 2011; Epp et al., 2012; Miya et al., 2015; Evans et al., 2016; Valentini et al., 2016, Berry et al., 2017; Ushio et al., 2018) and 16S (Taylor, 1996; Kitano et al., 2007; Evans et al., 2016; Vences et al., 2016; Bálint et al., 2018; Taberlet et al., 2018) rRNA primers for eDNA metabarcoding were then tested *in silico* with ecoPCR (Ficetola et al., 2010) against the available 12S and 16S sequences in the custom reference databases for vertebrate species of northern Alabama. During *in silico* testing, we allowed up to three mismatches between each reference sequence and each primer and 50–500 bp fragments. The 12S rRNA primers (12S-V5-F: 5′-ACTGGGATTAGATACCCC-3′ and 12S-V5-R: 5′-TAGAACAGGCTCCTCTAG-3′) developed by Riaz et al. (2011) amplified the greatest number of species *in silico* and thus we used these primers in this study. These primers amplify a 90-109 bp fragment of the 12S rRNA gene in Alabama vertebrates.

### 2 eDNA METABARCODING

#### 2.1 Library preparation

All metabarcoding (i.e., PCR and high-throughput sequencing) was conducted at the University of Illinois at Urbana-Champaign following clean protocols to minimize contamination (see Methods; e.g., frequent glove changes, use of UV on pipettes). We used a two-step PCR protocol following Harper et al. (2019) and Di Muri et al. (2020). Primers were modified to include molecular identifier (MID) tags, heterogeneity spacers, sequencing primers, and pre-adapters (giving a theoretical amplicon size of 243-269 bp). We ran PCRs in triplicate for each eDNA sample, with each plate using the same MID tags, following which replicates for each sample were pooled. Each sub-library contained at least two PCR negative controls (molecular grade water) and two PCR positive controls of greater prairie chicken (*Tympanuchus cupido*).

PCR1 reagents and conditions are described in Methods. To confirm successful amplification of eDNA samples and correct size of PCR products (~200–300 bp), we mixed 2 µL of PCR product with 0.5 µL of 5X DNA Loading Buffer Blue (Bioline®, Taunton, MA, USA) and visualized on a 2% gel. To make this gel, we mixed 70 mL 1X Sodium Borate buffer (Brody and Kern, 2004) with 1.4 g agarose powder (Fisher Scientific, Waltham, MA, USA) and GelRed to stain the gel (Biotium, Inc. Fremont, CA, USA). To enable size estimation, we used 1 µL of HyperLadder™ 100 bp (Bioline®, Taunton, MA, USA). We ran all gels at 200 V for 20 minutes.

After confirming successful amplification for the first PCR, we pooled PCR products by PCR plate into 16 sub-libraries based on band strength from gel images (Alberdi et al. 2018; see

Methods). We purified these sub-libraries using Mag-Bind® TotalPure NGS magnetic beads (Omega Bio-tek, USA) following a double size selection protocol (Bronner et al., 2009) at ratios of 0.9X and 0.15X magnetic beads to 100 µL of each sub-library to remove primer dimer (<200 bp) and secondary product (>1000 bp). We then visualized purified sub-libraries on a 2% agarose gel (following methods mentioned above) to confirm absence of primer dimer and secondary product. If either were present, we repeated bead clean-up on the affected sub-libraries and confirmed the lack of primer dimer or secondary product. We stored purified sub-libraries at -20°C until the second PCR.

PCR2 reagents and conditions are described in Methods. We pooled PCR replicates for each sub-library and visualized on a 2% agarose gel (as above) to confirm successful amplification. To remove secondary products, we again used a double size selection magnetic bead clean-up followed by confirmation of the absence of secondary bands on a 2% agarose gel (as above). Magnetic bead clean-up was performed with ratios of 0.7X and 0.15X Mag-Bind® TotalPure NGS magnetic beads to 50 µL of PCR product to remove primer dimer and secondary products but retain the target amplicon (~315 bp), confirmed by visualizing sub-libraries on a 2% agarose gel (as above). Sub-libraries were normalized based on number of samples they represented and estimated concentration using a Qubit™ dsDNA HS Assay Kit on a Qubit™ 3.0 fluorometer (Thermo Fisher Scientific™, Waltham, MA, US), then pooled. We performed a final magnetic bead clean-up (with 0.7X and 0.15X magnetic beads to 50 µL pooled library), diluted to 4 nM, and then sent to the Roy J. Carver Biotechnology Center, University of Illinois at Urbana-Champaign, for QA/QC and sequencing (see Methods).

### 2.2 Bioinformatics

We used a custom python script with metaBEAT v0.97.11 (https://github.com/HullUni-bioinformatics/metaBEAT) for bioinformatic processing and taxonomic assignment (Hänfling et al. 2016). Raw reads were quality trimmed from the read ends (minimum per base phred score Q30) and across sliding windows (window size 5 bp; minimum average phred score Q30) using Trimmomatic v0.32 (Bolger et al. 2014). The first 18 bp were removed to ensure no locus primer remained, reads were cropped to a maximum length of 114 bp, and any reads <90 bp were discarded. Then, we merged forward and reverse reads using FLASH v1.2.11 (Magoč and Salzberg, 2011), with a minimum of 10 bp overlap and a maximum of 10% mismatch between pairs. If reads did not merge, we retained only forward reads for maximum read quality. We used a final length filter to keep only sequences that were of the expected fragment size (90−110 bp). To remove chimeric sequences, we queried sequences against our custom 12S reference database using the *uchime* algorithm (Edgar et al., 2011) in vsearch (v1.1; Rognes et al., 2016). Also using vsearch, we clustered reads at 100% identity to remove any redundancy for taxonomic assignment. Clusters were considered sequencing error and omitted from further processing if they were represented by less than three sequences. Taxonomic assignment was performed with our custom 12S reference database using BLAST (Zhang et al., 2000). If any query sequence matched a reference sequence with at least 98% identity across more than 80% of its length, then taxonomic identity was assigned using a lowest common ancestor (LCA) approach based on the top 10% BLAST matches, similar to the strategy used by MEGAN (v5.10.6; Huson et al., 2007). Unassigned sequences were subjected to a separate BLAST search against the complete NCBI nucleotide database at 98% identity to determine the source via LCA as described above.

### 2.3 Contamination threshold

When manually examining the bioinformatics results and the sequence reads for each sample, we noticed a discrepancy with labeling, where high levels of reads assigned to our positive control, *Tympanuchus cupido,* were detected in an eDNA sample and a PCR negative control that were next to PCR positive controls that appeared to have failed. We believe for these four samples/controls (P7.NTC.001 and P7.PC.001 and Crowright-030 and P1.PC.002), we either pipetted the wrong DNA template into nearby wells (mixing up samples/controls as determined from plate layouts) or mixed-up sample labels during the input file of demultiplexing. More specifically, when examining the results, P7.PC.001 failed (no reads assigned to *Tympanuchus cupido* and only 1,976 reads identified to humans [*Homo sapiens*] were detected), whereas nearby control P7.NTC.001 had 153,622 reads assigned to *T. cupido* (and 138 reads assigned to Phasianidae, the pheasant family) and 65 reads assigned to white-footed mouse (*Peromyscus leucopus*; contamination in our positive control addressed below). Similarly, sample Crowright-030 had 56,125 reads assigned to *T. cupido* (51 reads were assigned to Phasianidae), while P1.PC.002 did not have any reads assigned to *T. cupido* (but did have 794 reads assigned to redspotted sunfish, [*Lepomis miniatus]*, 66 reads assigned to Centrarchidae [sunfish family], and 480 reads assigned Percidae [perch family]). We corrected the labeling of these samples (P7.NTC.001 was switched with P7.PC.001; Crowright-030 was switched with P1.PC.002).

Initially, we evaluated several types of false positive sequence threshold commonly used in the eDNA metabarcoding literature (Drake et al. 2022), including 1) the maximum frequency of reads in the negative process controls (field blanks, extraction blanks, and NTCs), 2) the maximum frequency of positive control DNA in eDNA samples, 3) the maximum frequency of reads that were not *T. cupido* in the PCR positive controls, and 4) the maximum frequency of reads for each taxon in all process controls. These thresholds were applied to the data along with blank correction (see Methods) to assess their impact on contamination and plausible detections. See Figure 1 below for the level of contamination across our controls (field blanks, extraction blanks, NTCs, and our positive control).

For option 1, reads in negative controls were largely *H. sapiens*, domestic species (e.g., cattle [*Bos taurus*], chicken [*Gallus gallus*]), or taxa that could have been positive control DNA (e.g. Phasianidae). Wild taxa occurring at the next highest frequencies were stoneroller (*Campostoma*; 1.72%), darters (*Etheostoma*; 1.5%), and North American beaver (*Castor canadensis*; 1.02%). For option 2, positive control DNA was detected in nine eDNA samples at a range of frequencies (0.006-19.96%). For option 3, *P. leucopus* was the most severe contaminant at 0.042% after reads which could be positive control DNA. For option 4, there were 25 taxon-specific sequence thresholds ranging from 0.0009-85.35% (excluding *H. sapiens*, *G. gallus*, and *T. cupido* which were 100%). Options 1 and 2 would result in too many plausible detections being lost and option 4 is a less commonly used sequence threshold. This led us to choose option 3 but given that no fish reads were detected in PCR positive controls, we opted to use a fish threshold of 0% and a non-fish threshold of 0.042% with blank correction. This would account for contamination whilst retaining plausible low density and rare species that would be important to IBI calculations and maximize the total number of fish species detected. We then retained all taxonomic assignments with ≥ 3 sequence reads (noise threshold) to account for potential PCR or sequencing artefacts whilst retaining rare species that may be underrepresented in sequencing. Threshold evaluation and blank correction for contamination were performed using and before/after figures (see Figure 2 below) created using RStudio (v.4.4.1; R Core Team, 2024) with *plyr* (Wickham, 2011), *tidyverse* (Wickham et al., 2019), *ggpubr* (Kassambara, 2023), *scales* (Wickham et al., 2023), and *reshape2* (Wickham, 2007) packages.

### 4. Figures

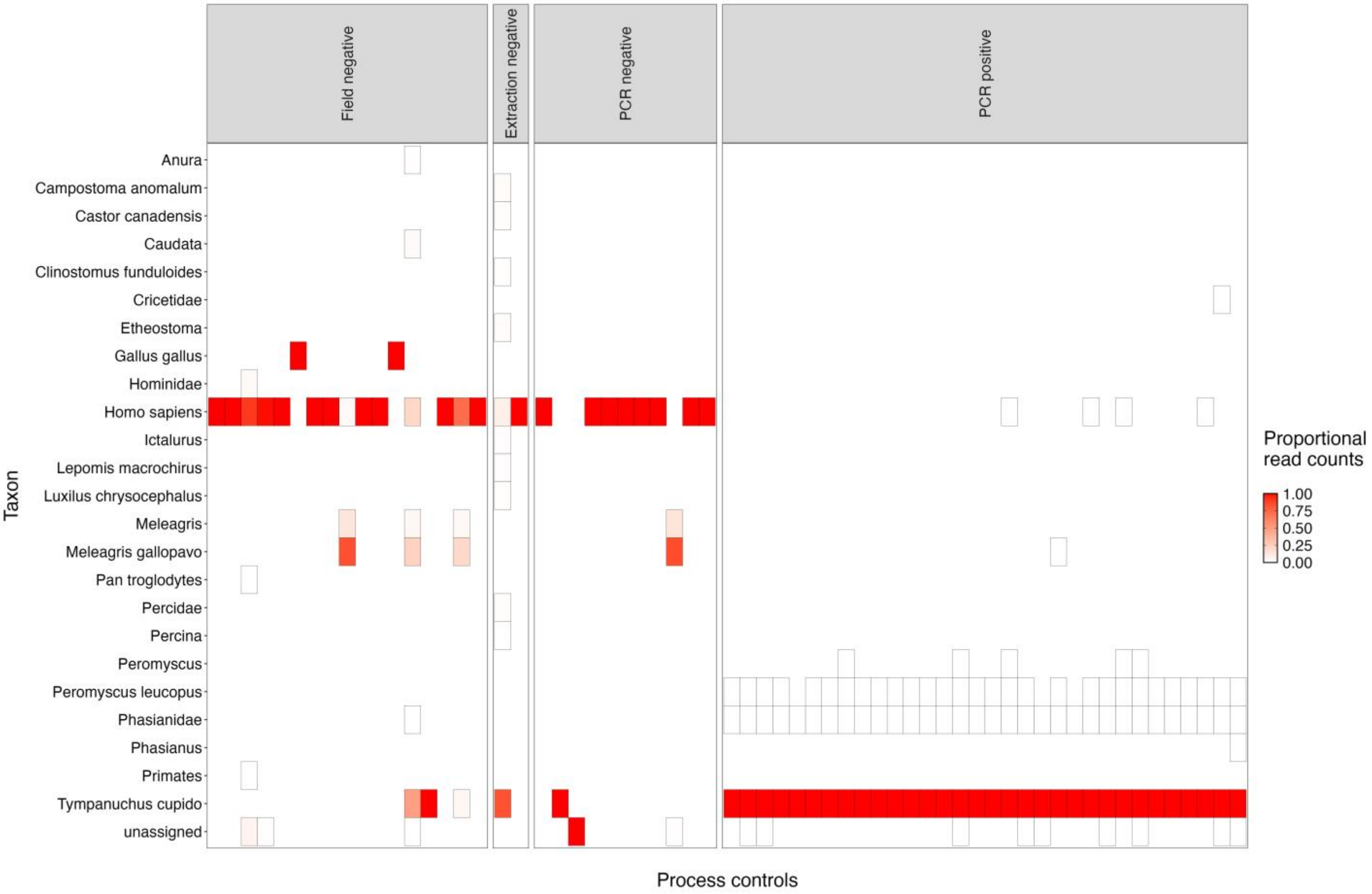


**FIGURE 1** The level (as proportional read counts) of contamination (by taxon) across controls/blanks (field negatives, extraction negatives, PCR negatives [NTC]), and PCR positives (extracted *T. cupio* DNA was used as our positive control). We used this information to assess different contamination thresholds.

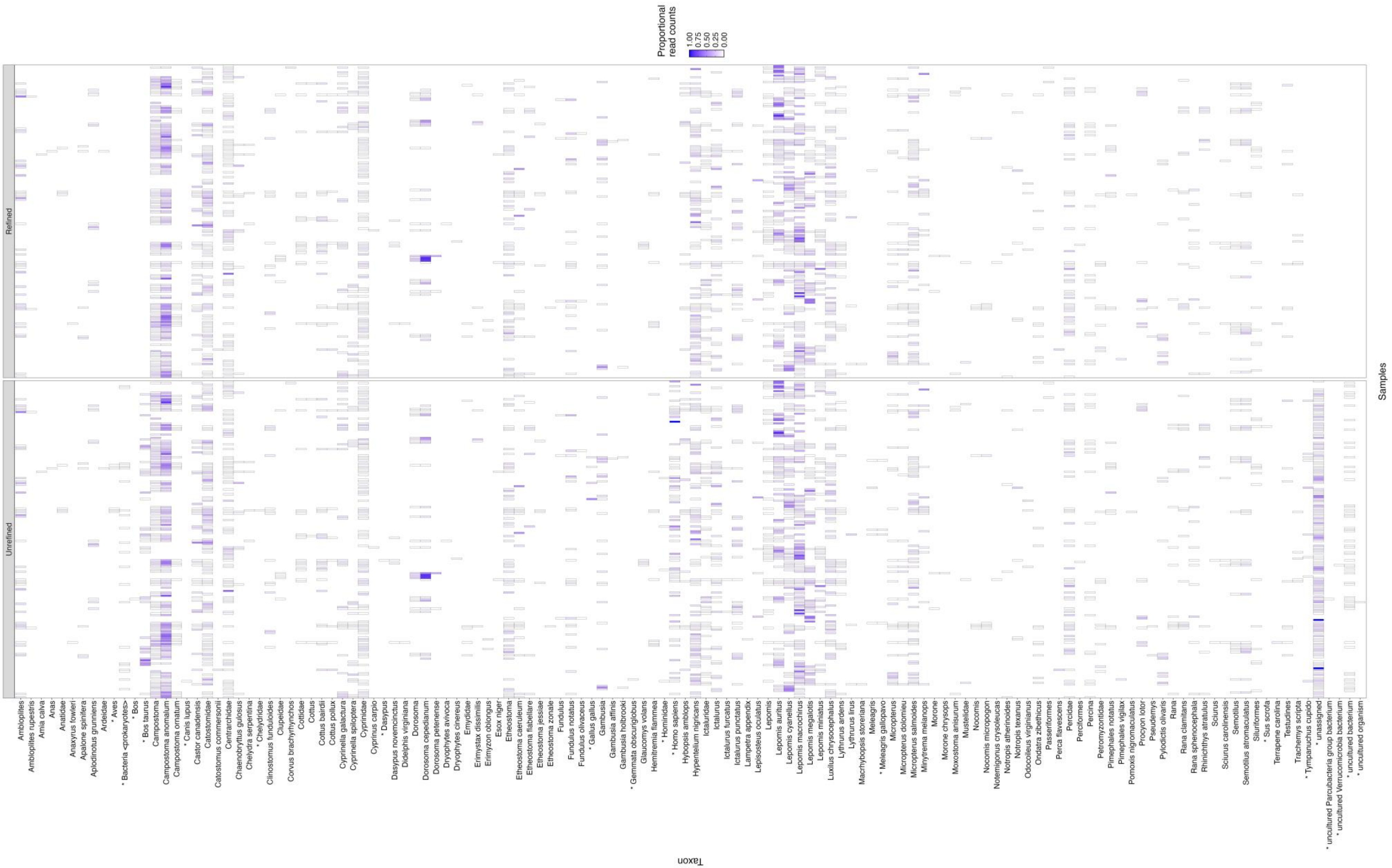


**FIGURE 2** The before (unrefined) and after (refined) of all taxon detected in our eDNA samples after we applied our contamination threshold (fish threshold or Option 3 in-text).
